# Co-occurrence of contrasting life-history strategies in a metapopulation inhabiting temporally variable and stable breeding sites

**DOI:** 10.1101/500942

**Authors:** Hugo Cayuela, Sam S. Cruickshank, Hannelore Brandt, Arpat Ozgul, Benedikt R. Schmidt

**Affiliations:** Institut de Biologie Intégrative et des Systèmes (IBIS), Université Laval, Québec, QC G1V 0A6, Canada; Institut für Evolutionsbiologie und Umweltwissenschaften, Universität Zürich, Winterthurerstrasse 190, 8057 Zürich, Switzerland; Swiss Federal Institute for Forest, Snow and Landscape Research (WSL), Zürcherstrasse 111, 8903 Birmensdorf, Switzerland; Info fauna karch, UniMail, Bâtiment G, Bellevaux 51, 2000 Neuchâtel, Switzerland

**Keywords:** amphibian, demography, life history, intraspecific variation, habitat stability

## Abstract

Life-history theory states that, during the lifetime of an individual, resources are allocated to either somatic maintenance or reproduction. Resource allocation trade-offs determine the evolution and ecology of life-history strategies and determine an organisms’ position along the fast-slow continuum. Theory predicts that environmental stochasticity is an important driver of resource allocation and therefore life-history evolution. Highly stochastic environments are expected to increase uncertainty in reproductive success and select for iteroparity and a slowing down of the life history. To date, most empirical studies have used comparisons among species to examine these theoretical predictions. By contrast, few have investigated how environmental stochasticity affects life-history strategies at the intraspecific level. In this study, we examined how variation in breeding site stochasticity (among-year variability in pond volume and hydroperiod) promotes the co-occurrence of different life-history strategies in a spatially structured population, and determines life-history position along the fast-slow continuum in the yellow-bellied toad (*Bombina variegata*). We collected mark-recapture data from a metapopulation and used multievent capture-recapture models to estimate survival, recruitment and breeding probabilities. We found higher survival and longer lifespans in populations inhabiting variable sites compared to those breeding in stable ones. In addition, probabilities of recruitment and skipping a breeding event were higher in variable sites. The temporal variance of survival and recruitment probabilities as well as the probability to skip breeding was higher in variable sites. Taken together, these findings indicate that populations breeding in variable sites experienced a slowing down of the life-history. Our study thus revealed similarities in the macroevolutionary and microevolutionary processes shaping life-history evolution.

## Introduction

Population biologists strive to understand the ecological mechanisms that drive life-history evolution. Life-history theory is based on the premise that during the lifetime of an individual, energy and resources are allocated to either somatic maintenance, growth, or reproduction (Stearns 1976, 1992, 2000). Life-history tradeoffs occur when a beneficial change for one of these traits entails a detrimental change for another (Stearns 1976, 1992, 1989). For example, resources which are invested into reproduction cannot be used for somatic maintenance or growth, which may lead to shorter lifespans. Trade-offs between traits shape the architecture of life-history strategies and the distribution of organisms along the fast-slow continuum (Sæther & Bakke 2000, Sæther et al. 2004, Salguero-Gómez et al. 2016). At the fast end of the continuum, organisms have rapid growth, short lifespans and high fecundity (large broods and little parental care); at the slow end, they have opposite characteristics (Stearns 1983). In “slow” species, natural selection favors the evolution of an environmental canalization process which reduces annual variation in adult survival (Gaillard & Yoccoz 2003). Indeed, adult survival is the life-history trait which has largest impact on population growth rate in long-lived iteroparous organisms (Pfister 1998, Gaillard & Yoccoz 2003, Hamel et al. 2010).

Over the last decades, theoretical studies have emphasized the central role of environmental stochasticity in life-history evolution (Gillespie 1977, Bulmer 1985, Tuljapurkar 2013). Reproductive success is often uncertain in stochastic environments. In amphibians, for example, weather conditions determine pond hydroperiod and pond hydroperiod determines survival of larvae (Pechmann et al. 1989). Variation in reproductive success due to environmental stochasticity can select for iteroparity and a slowing down of the life history (Bulmer 1985, Tuljapurkar et al. 2009, Tuljapurkar 2013). Iteroparity allows individuals to exploit multiple breeding occasions, which mitigates the risk of reproductive failure when environmental conditions are unfavorable in some years. In addition, reproductive effort can be phenotypically plastic in iteroparous species. The mechanisms underlying plasticity in reproductive effort can be behavioural or physiological. Individuals may decide not to breed in unfavourable years and skip opportunities for breeding (Bull and Shine 1979, Pilastro et al. 2003, Skjæraasen et al. 2012). Individuals may also adjust clutch size (Boyce & Perrins 1987, Baker et al. 2015) or resorb oocytes (Moore & Attisano 2011, Attisano et al. 2013).

To date, most of the empirical studies that have examined the effects of environmental stochasticity upon life-history variation have focused their attention at the interspecific level (e.g. Bielby et al. 2007, Forcada et al. 2008, Morris et al. 2008). By contrast, very few studies have tested theoretical predictions at the intraspecific level (e.g., Semlitsch et al. 1990, Nevoux et al. 2010), probably due to the scarcity of individual data collected in wild populations experiencing contrasting regimes of environmental stochasticity. In one study, Nevoux et al. (2010) compared the life-history strategy of two bird populations and showed that an increased climatic variability in the Atlantic Ocean resulted in a longer lifespan and lower reproductive success compared to another population in the Indian Ocean. Here, we aimed at examining how spatial variation in environmental stochasticity leads to the co-occurrence of contrasting demographic strategies within the same metapopulation, and determines population position along the fast-slow continuum.

Pond-breeding amphibians may serve as informative biological models to deepen our understanding of these issues. Many species reproduce in ponds where reproductive success (i.e., larval survival) varies among years. Much of this variability is caused by variation in pond hydroperiod over space and time (Shoop 1974, Pechmann et al. 1989, McCaffery et al. 2014). Moreover, amphibians can display broad intraspecific variation in their demographic traits (i.e., survival and fecundity; Berven & Gill 1983; Morrison & Hero 2003; Cayuela et al. 2017, Muths et al. 2017). Reproductive effort is highly flexible in amphibians because individuals can skip opportunities for reproduction (Husting 1965, Church et al. 2007, Muths et al. 2006, Cayuela et al. 2016b), adjust clutch and egg size (Crump 1981, Reyer et al. 1999) and resorb oocytes (Jørgensen 1984) according to environmental conditions.

In this study, we examined how spatial variation in breeding pond stochasticity affects population position along the fast-slow life-history continuum in a metapopulation of yellow-bellied toads (*Bombina variegata*). *B. variegata* is a prolonged breeder (*sensu* Wells 1977). The long breeding season usually begins in May and lasts until July (Barandun and Reyer 1997). Females can lay multiple clutches in a season (Buschmann 1998). We collected capture-recapture data in a metapopulation composed of 14 populations during a 5-year period. In the metapopulation that we studied, populations inhabit two contrasting types of breeding site (Fig. 1). Two populations reproduce in ponds whose the water level varies dramatically throughout the breeding season (hereafter called “variable breeding sites”). Ponds are dry most of the season and fill after heavy rainfall (Fig. 1). Depending on rainfall, pond volumes are small or large and hydroperiods can be short or long (Pechmann et al. 1991). The other populations live in habitats where the fluctuations in the water level and volume of the ponds are small. It is rare that ponds dry and the areas are almost never flooded (hereafter “stable breeding sites”; Fig.1) Notably, the stable ponds are man-made ponds. Differences in pond volume and length of hydroperiod are well known to affect freshwater organisms including amphibians through effects upon abiotic conditions (e.g., bathymetry) and biotic interactions (e.g., predators; Pearman 1993, Wellborn et al. 1996, Adams et al. 2003, McCaffery et al. 2014).

**Fig. 1.**
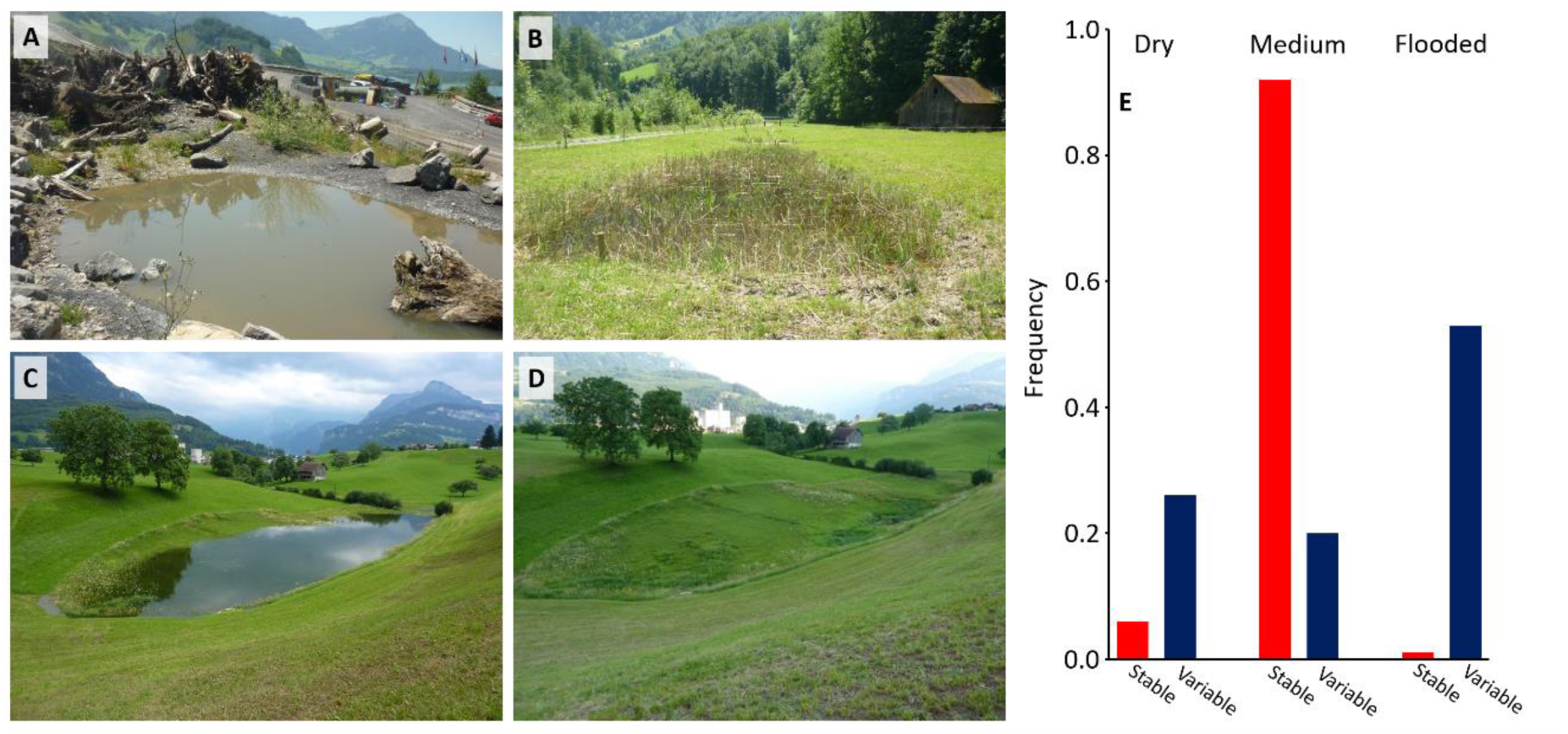
Pictures of stable and variable breeding sites in the study area. The pictures A and B show two stable sites (“Steinbruch Zingel”, A; “Hinter Ibach”, B) where water levels do not vary much. The two pictures C and D show the variable site “Wilen” where the pond is either dry or the the site is flooded, depending on rainfall. (E) Description of water levels of ponds in stable and variable sites. At every capture occasion, we assessed qualitatively pond water levels. The figure shows the proportion of capture occasion in which ponds in a site were found in one of the three states (dry, medium, flooded).

Using multievent capture-recapture models (Pradel 2005), we tested a set of hypotheses. In *B. variegata*, adult survival has been reported to be the demographic parameter having the largest effect on population growth rate (Cayuela et al. 2015). Therefore, we hypothesized that (1) adult survival probability would have a lower temporal variance than recruitment probability (regardless of site type), indicating a higher environmental canalization (i.e., buffering) of adult survival (Pfister 1998, Gaillard & Yoccoz 2003). According to life-history theory (Stearns 1976, Gillespie 1977, Bulmer 1985, Tuljapurkar 2013), we expected (2) higher survival (and therefore longer lifespan) in populations reproducing in more variable sites compared to populations breeding in less variable ones. (3) Amphibians are well-known to skip breeding in years when conditions are not favourable (Church et al. 2007, Muths et al. 2006). We hypothesized that the probability of skipping breeding would be higher and more variable through time in populations where breeding sites were variable, when compared with populations breeding in stable sites. (4) Moreover, we formulated two alternative hypotheses about the recruitment of first breeders into the population: recruitment rate would be lower and more variable in variable sites, possibly as a result of lower and more variable breeding effort (hypothesis 3); (5) alternatively, recruitment rate would be higher and more variable in variable sites, possibly due to a breeding effort synchronized with environmental variation (and positively affecting breeding success). Finally, according to life-history theory (life-history evolution (Gillespie 1977, Bulmer 1985, Tuljapurkar 2013), we formulated the general prediction that (6) populations reproducing in variable sites would experience a life-history slowdown.

## Materials and methods

### Study area, site classification and capture-recapture survey

The study area is situated within the Schwyz-Ingenbohl basin in central Switzerland. The data used in this study come from a monitoring program for a metapopulation of *B.variegata* that was set up to evaluate the effects of pond-building on the species. The valley basin was comprehensively surveyed prior to the start of the capture-recapture study. There are 14 sites with *variegata* populations (T. Hertach and M. Schlitner, personal communication). Sites consisted of between 1 and 15 ponds situated in close proximity to one another. The majority of sites were situated within agricultural meadows or within active quarries. We classified two of the sites as “variable” breeding sites as the water levels and pond volumes of the sites varied dramatically throughout the breeding season caused strong variation among years in the length of the hydroperiod (Fig. 1). All other sites were classified as “stable” as pond water levels remained relatively stable throughout the breeding season.

Toads were surveyed using the capture-recapture method over a 5-year period (2011-2015). Captures were carried out during the breeding season (May-August), with between two and four capture events per site per year; the number of individuals captured at each session in variable and stable sites is given in Supplementary material 1, Table S1. Toads were captured by hand during the hours of night (21:00-04:00), and individuals were photographed to allow identification from the ventral patterns of black and yellow mottles (Cruickshank & Schmidt 2017). We are confident that the survival estimates are not negatively biased by permanent emigration as we exhaustively surveyed all the populations comprising the metapopulation and because we detected very few dispersing individuals (11 out of 3822 known individuals).

### Modeling survival and breeding probabilities

We used multievent models (Pradel 2005) to estimate survival, recruitment, breeding, and recapture probabilities. An event is what is observed in the field and thus coded in the individual capture history. This observation is related to the latent state of the individuals. Yet, observations can come with a certain degree of uncertainty regarding the latent state. Multievent models aim at modelling this uncertainty in the observation process using hidden Markov chains.

The models use the robust-design structure (Pollock 1982) to quantify changes in individuals breeding status. In amphibians, robust-design models have been used to quantify breeding probability and its complement, the probability of skipping a breeding event (Bailey et al. 2004, Muths et al. 2006; Church et al. 2007, Cayuela et al. 2014). Breeding amphibians are usually captured in aquatic sites. Non-breeding amphibians located in terrestrial environments are by definition not available for capture. If an individual does not breed at a given occasion, it skips a breeding opportunity. Technically, skipping breeding is equivalent to becoming a temporary emigrant and moving to an unobservable state (Kendall & Nichols 2002, Schaub et al. 2004). In our model, we estimated both early and late season probabilities of skipping breeding. The former corresponds to the case when a toad skips skipping at the first capture occasion of the year. Late season probabilities of skipping a breeding opportunity describe the probability that a toad is absent from the breeding site later in the season in a year (i.e., during capture occasion 2 to 4 in year *i*). The model also included heterogeneity mixtures (Pledger 2000, Gimenez & Choquet 2010) to account for inter-individual heterogeneity in recapture probabilities. Heterogeneity models assign individuals to one of two groups where one group has a higher detection probability.

The multievent model incorporated 5 composite states (B1, B2, NB1, NB2 and the dead state D) that include information about the breeding status of individuals (“B” for breeders and “NB” for non-breeder) and heterogeneity group. Two heterogeneity groups are considered: ‘1’ for individuals in the group which are easily captured, and ‘2’ for ones in the group which are harder to capture. Two potential events were considered in the model: captured (“C”) or not (“NC”) during a given occasion. At their first capture, individuals can be in either state B1 or state B2 (Fig. 2). Note that individuals cannot be in a non-breeding state at their first capture as non-breeding individuals are, by definition, unavailable for capture. To model state–state transitions between times *t*-1 and *t*, we consider two successive steps during which the information of the state descriptor is progressively updated (Pradel 2005): (1) survival and (2) breeding probability (Fig. 2 and 3). Each step is conditional on all previous steps. States and transitions are summarized in a transition matrix (Fig. 2). The rows of the matrix correspond to time *t*-1 and the columns to time *t*.

**Fig. 2.**
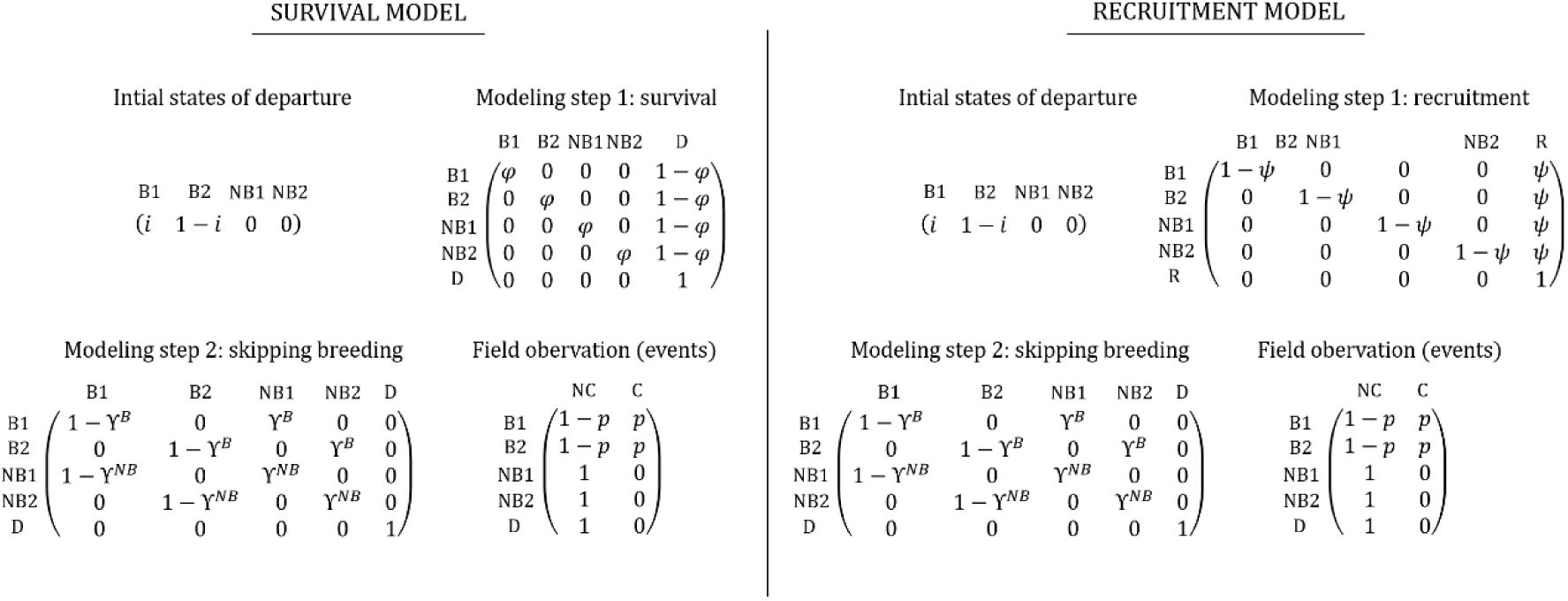
Survival and recruitment analyses: structure of the multievent models.

**Fig. 3.**
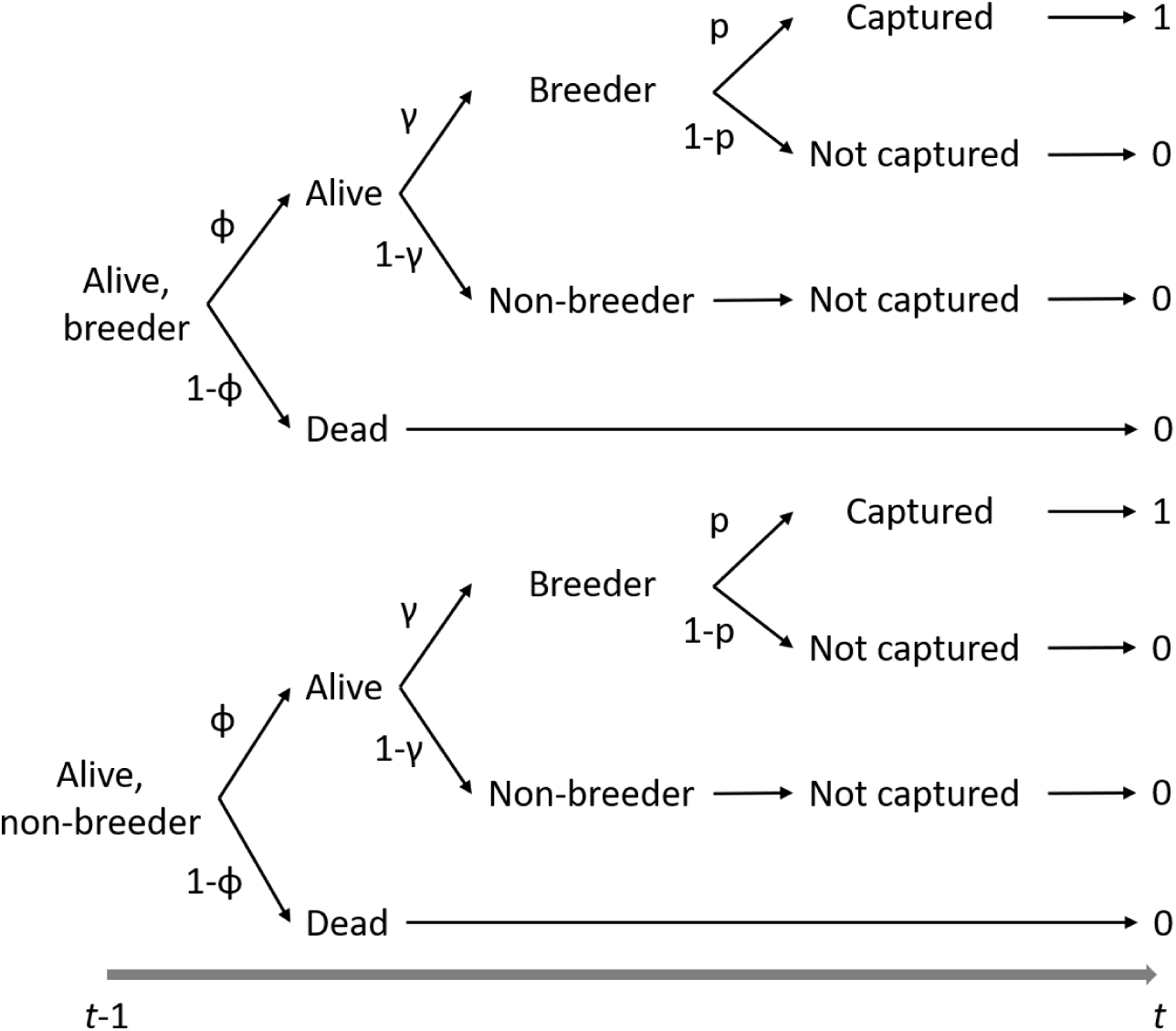
Diagram of fate of an adult of Bombina variegata surveyed in the metapopulation between two capture sessions t-1 and t. The transition probabilities correspond to the annual survival (φ) and the probability of skipping breeding (γ). The event probabilities correspond to the probabilities of being captured (p) in breeding sites.

In the first modeling step, survival information is updated and an individual can survive with a probability φ or may die with a probability 1-φ; if it dies, it enters state D and remains in this state. This leads in a transition matrix with 5 states of departure and 5 states of arrival (Fig. 2). In the second modeling step, the information about individual breeding status (B or NB) is updated. An individual that has survived can breed or skip a breeding opportunity at *t* with probability 1-γ^*B*^ and γ^*B*^, respectively, given it has bred at *t*-1. An individual that has already skipped breeding at *t*-1 may skip a breeding occasion at *t* with a probability γ^*NB*^. This results in a transition matrix with 5 states of departure and 5 states of arrival (Fig. 2). The last component of the model links field observation (i.e., events) to these underlying true states. Breeding individuals can be recaptured with probability *p*. Non-breeding and dead individuals cannot be recaptured, hence their recapture probability *p* is fixed at zero.

### Modeling recruitment and skipping breeding behavior

We examined recruitment using a model following the structure of Pradel’s (1996) model. In our modified version of the Pradel model (Fig. 2), recruitment probability was estimated as the probability that an individual present at t was not present at t-1, i.e. the proportion of “new” individuals in the population at *t*, while accounting for recapture heterogeneity. The model included the same five states B1, B2, NB1, NB2 and R (replacing D), as in the model described above. The state R correspond to “being recruited”. The steps of initial states, the breeder-non/breeder transitions and the events were similar to those of the previous model (Fig. 2). However, we introduced a recruitment step (replacing the survival step) in which recruitment information was updated; between t-1 and t, an individual could be recruited with a probability Ψ. This leads to a transition matrix with 5 states of departure and 5 states of arrival (Fig. 2).

### Building biological scenarios

The survival and recruitment models were implemented in program E-SURGE (Choquet et al. 2009). We ranked models using the Akaike information criterion adjusted for a small sample size (AICc) and Akaike weights (w). If the Akaike weight of the best supported model was less 0.9, we used model-averaging to obtain parameter estimates (Burnham & Anderson 2002). The 95% CI were calculated using the delta-method (Royall 1986). We tested our hypotheses about survival, the probabilities of recapture, and to skip breeding, from the following general model: [φ(Habitat), γ_*late*_(Habitat), γ_*early*_(Habitat), p(Het + Year + Habitat)]. Model notation follows the standard notation of Lebreton et al. (1992). The variables in the models were site type (Habitat: stable vs variable), heterogeneity mixture (Het) and year (Year). We hypothesized that: (1) survival φ probability and skipped breeding probabilities both early (γ_*inter*_) and late (γ_*intra*_) in the season differed between stable sites and variable sites (Habitat). The site effect was included in the model via a discrete covariable with two modalities (i.e. “stable” vs “variable”). Concerning the probability to skip breeding, we considered the possibility that individuals may leave or enter into breeding ponds either early or late in the season. To avoid identifiability issues, we assumed that the probability to skip breeding did not depend on individuals’ past breeding status; consequently, the probabilities γ^*B*^ and γ^*NB*^ were set equal in the models. Recapture probability p was assumed to depend on two heterogeneity groups (Het). We also hypothesized that recapture probability varied between years (Year) and according to site stability/variability (Habitat). We tested all possible combinations of effects, resulting in the consideration of 32 competing models (Table 1). Moreover, we expected that skipping behavior should not differ between sexes with respect to water availability; the absence of water in ponds will result in lack of breeding attendance in both sexes.

**Table 1.** Survival analyses: model selection results. We show the 5 best-supported models based on their AICc rank (the complete selection procedure is provided in Supplementary material 1). The models include four parameters: φ **=** survival, γ_late_ = late season skipping breeding probability, γ_early_ = early season skipping breeding probability, p = recapture probability. k = number of model parameters, Deviance = residual deviance, ΔAICc = difference of AICc points between the model and the best-supported model, w = Akaike weight. The variables in the models were site type (stable vs variable, Habitat), heterogeneity mixtures (low vs high recapture probability, Het) and year (Year).

| Model | k | Deviance | $\Delta AICc$ | w |
| --- | --- | --- | --- | --- |
| $\phi(\text{Habitat}), Y_{early}(\text{Habitat}), Y_{late}(\text{Habitat}), p(\text{Het+Year+Habitat})$ | 14 | 16772.94 | 0.00 | 0.57 |
| $\phi(\text{Habitat}), Y_{early}(\text{Habitat}), Y_{late}(\text{Habitat}), p(\text{Het+Year})$ | 13 | 16775.58 | 0.63 | 0.42 |
| $\phi(\cdot), Y_{early}(\text{Habitat}), Y_{late}(\text{Habitat}), p(\text{Het+Year+Habitat})$ | 13 | 16783.35 | 8.40 | 0.01 |
| $\phi(\text{Habitat}), Y_{early}(\cdot), Y_{late}(\cdot), p(\text{Het+Year+Habitat})$ | 13 | 16786.71 | 11.76 | 0.00 |
| $\phi(\cdot), Y_{early}(\text{Habitat}), Y_{late}(\text{Habitat}), p(\text{Het+Year})$ | 12 | 16790.66 | 13.70 | 0.00 |

We tested our hypotheses on recruitment using the following general model [Ψ(Habitat), γ_*late*_(Habitat), γ_*early*_(Habitat), p(Het + Year + Habitat)]. In this model, we hypothesized that recruitment (Ψ) and probabilities of skipping breeding (γ) differed between variable and stable sites. We also assumed that recapture probability depended on two heterogeneity groups and varied between years and according to site stability/variability. All the possible combinations of effects were considered, leading to 32 candidate models (Table 1).

### Estimating temporal variance of survival, breeding and recruitment probabilities

We ran a model with annual variation in survival and the probabilities of skipping breeding [φ(Habitat × Year), γ_*late*_(Habitat × Year), γ_*early*_(Habitat × Year), p(Het + Year + Habitat)] to obtain annual estimates of survival and probabilities of skipping breeding, along with their estimated variance and covariance. Using the methods proposed in Gould and Nichols (1998), we estimated the temporal variance in survival and the probabilities of skipping breeding by removing the effects of sampling variance from the estimated total variance. As in Cayuela et al. (2016), we used these calculated standard deviations as a proxy for estimating the presence of environmental canalization (Gaillard & Yoccoz 2003) of survival and breeding probabilities. We then used the method proposed by Gaillard & Yoccoz (2003) to compare the temporal variation in the variance of survival probabilities and probabilities of skipping breeding. This method compares the maximum variance we can expect with the observed variance in demographic rates. For instance, the maximum variance of survival is φ × (1 - φ), where φ is the mean survival rate. Between the observed variance and the maximum variance, we then obtained “temporal variation ratio” (following the terminology used by Gaillard & Yoccoz 2003), which can be compared between demographic rates and sites. This approach was used to quantify temporal variation ratio of recruitment, from the model [Ψ(Habitat × Year), γ_*late*_(Habitat × Year), γ_*early*_(Habitat × Year), p(Het + Year + Habitat)].

## Results

### Survival, probability to skip breeding and recruitment in stable and variable sites

The best-supported model for survival was the general model [φ(Habitat), γ_*late*_(Habitat), γ_*early*_(Habitat), *p*(Het + Year + Habitat)] (Table 1). Yet, as its Akaike weight was 0.89, we used model-averaging to obtain parameter estimates. Recapture probability varied according to heterogeneity group, year and site stability/variability (see Supplementary material 1, Fig. S1). The recapture probability of the individuals assigned to the heterogeneity mixture 1 varied between 0.40 (95% CI 0.31-0.49) in 2013 and 0.98 (95% CI 0.87-0.99) in 2011 while it ranged from 0.02 (95% CI 0.01-0.05) in 2013 to 0.55 (95% CI 0.32-0.76) in 2011 in individuals assigned to the heterogeneity mixture 2. The recapture probability was marginally higher in variable sites than in stable ones (Supplementary material 1, Fig. S1).

Survival was higher in variable sites than in stable ones (Fig. 4A). In variable sites, survival was 0.85 (95% CI 0.75-0.92) while it was 0.71 (95% CI 0.68-0.74) stable sites. This difference in annual survival results in a life expectancy of 5.67±2.42 and 2.45±0.90 years in variable and stable sites, respectively (life expectancy was calculated using the expression survival/(1-survival)). In variable sites, individuals also had a higher probability to skip breeding than in stable ones (Fig. 4C-D). Early in the season, an individual from a variable site had a probability to skip breeding of 0.76 (95% CI 0.69-0.84) whereas it was 0.62 (95% CI 0.53-0.69) in stable sites. Later in the season, the probability that an individual skips a breeding opportunity was 0.72 (95% CI 0.63-0.79) in variable sites while it was 0.56 (95% CI 0.47-0.64) in stable sites.

**Fig. 4.**
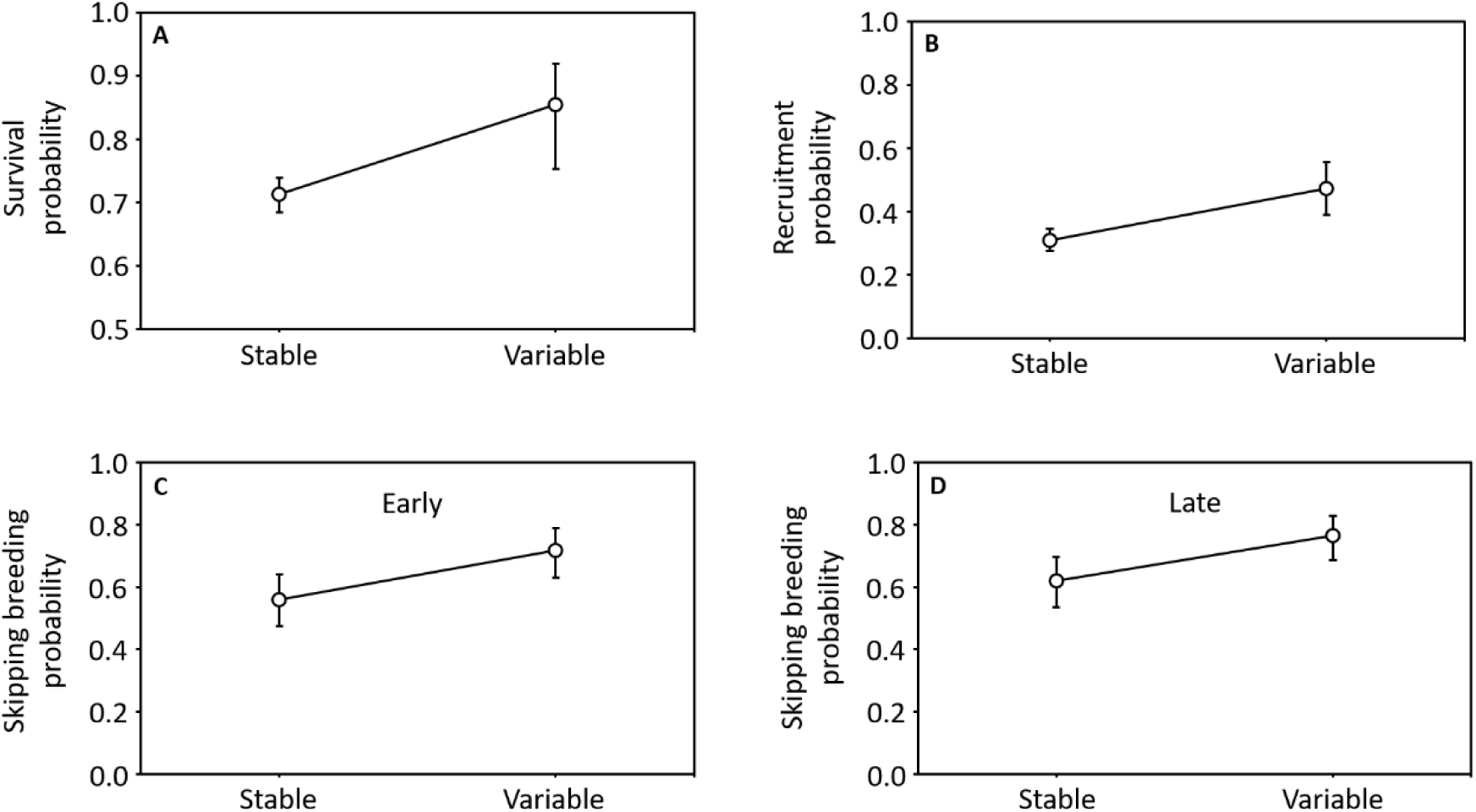
Influence of site stability/variability on survival (A), recruitment (B) and skipping breeding (C and D) probabilities (early and late). Estimates were model-averaged (from the models whose the sum of the AICc weight was 0.99; Table 1 and 2) and 95% CI (error bars) were calculated using the delta-method.

The best-supported model for recruitment was the general model [Ψ(Habitat), γ_*late*_(Habitat), γ_*early*_(Habitat), *p*(Het + Year + Habitat)]. Our results revealed that recruitment probability was higher in variable sites than in stable ones: in the former, it was 0.47 (95% CI 0.39-0.55) while it was 0.31 (95% CI 0.27-0.34) in the latter.

**Table 2.** Recruitment analyses: model selection results. We show the 5 best-supported models based on their AICc rank (the complete selection procedure is provided in Supplementary material 1). The models include four parameters: Ψ **=** recruitment, γ_late_ = late season skipping breeding probability, γ_early_ = early season skipping breeding probability, p = recapture probability. k = number of model parameters, Deviance = residual deviance, ΔAICc = difference of AICc points between the model and the best-supported model, w = Akaike weight.

| Model | k | Deviance | $\Delta AICc$ | w |
| --- | --- | --- | --- | --- |
| $\psi(\text{Habitat}), Y_{early}(\text{Habitat}), Y_{late}(\text{Habitat}), p(\text{Het+Year+Habitat})$ | 14 | 18098.58 | 0.00 | 0.89 |
| $\psi(\cdot), Y_{early}(\text{Habitat}), Y_{late}(\text{Habitat}), p(\text{Het+Year+Habitat})$ | 13 | 18104.85 | 4.25 | 0.11 |
| $\psi(\cdot), Y_{early}(\text{Habitat}), Y_{late}(\text{Habitat}), p(\text{Het+Year})$ | 12 | 18113.02 | 10.42 | 0.00 |
| $\psi(\text{Habitat}), Y_{early}(\text{Habitat}), Y_{late}(\text{Habitat}), p(\text{Het+Year})$ | 13 | 18112.79 | 12.20 | 0.00 |
| $\psi(\cdot), Y_{early}(\cdot), Y_{late}(\cdot), p(\text{Het+Year+Habitat})$ | 12 | 18139.94 | 37.34 | 0.00 |

### Temporal variances of survival, probability to skip breeding and recruitment in stable and variable sites

The temporal variation ratios indicate that survival (0.07 and 0.16 in stable and variable sites, respectively) was less variable than recruitment (0.36 and 0.47 in stable and variable sites, respectively). Our analyses also showed that all the demographic parameters were more variable in variable sites than in stable ones. Temporal ratio of survival was 0.16 in variable sites whereas it was 0.07 in stable ones. The probabilities of skipping breeding were also more temporally variable in variable sites (early: 0.33, late= 0.09) than in stable ones (early: 0.17, late: 0.02). Moreover, temporal ratio for recruitment was 0.47 in variable sites and 0.36 in stable ones.

## Discussion

Our study revealed that adult survival probability had a lower temporal variance than recruitment probability, indicating that adult survival was more canalized than recruitment (confirming hypothesis 1). The results show higher survival and longer lifespan in populations reproducing in variable sites compared to populations breeding in stable ones (hypothesis 2). Furthermore, the analysis showed that the probabilities of skipping breeding were higher and more variable in populations in variable sites (hypothesis 3). In addition, we showed that recruitment probability was higher and more variable in variable sites (hypothesis 5). Overall, our study showed that populations breeding in variable sites experienced a slowing down of their life history (hypothesis 6).

### Stability and variability of breeding sites affects adult survival

Adult survival had a lower temporal variance ratio than recruitment in both stable and variable sites, indicating a higher level of canalization of adult survival. This result supports the conclusion of previous studies that adult survival is the most canalized demographic parameter in this species due to high sensitivity of population growth rate to variation in adult survival (Cayuela et al. 2015, 2016c). The analysis also revealed that adult survival depended on the level of breeding site stability. Survival was 16% lower in stable sites (φ = 0.71) compared to variable ones (φ = 0.85). This difference is substantial in light of the range of variation in adult survival (minimum: 0.71, maximum: 0.92) reported in *B. variegata* populations from western Europe (Cayuela et al. 2016a). In our study system, variation in adult survival was associated with changes in breeding effort (discussed below), and annual survival probability positively covaried with the probability to skip breeding in the populations. By skipping breeding, most of the costs of breeding are likely avoided, and the saved energy could be possibly re-allocated to somatic maintenance and subsequent reproduction. Indeed, female amphibians have the ability to resorb oocytes when skipping reproduction (Guarino et al. 1998; Reyer et al. 1999). Males can also save the energy not invested in activities associated with reproduction (e.g. calling and territorial behavior) that are highly energy consuming in amphibians (Wells 1977, Grafe & Thein 2001, Ophir et al. 2010). Therefore, it seems very likely that a decrease in breeding effort in variable sites contributes to the increase in adult survival and the doubling of the expected lifespan (5.6 years in variable sites vs 2.4 years in stable sites).

### Stability and variability of breeding sites affects breeding effort and recruitment

Our study showed that the probability of skipping breeding was higher and temporally more variable in variable sites than in stable ones. These results are line with the conclusions of previous studies, and show that amphibians may adjust their reproductive effort according to external factors (Church et al. 2007, Walls et al. 2013, Ficetola et al. 2016). Among the external factors, pond hydroperiod is particularly important since it has a strong influence on breeding success (Pechmann et al. 1989, Richter et al. 2003, Taylor et al. 2006, McCaffery et al. 2014). This is also congruent with our knowledge about natural history of *B. variegata*. Previous studies have shown that breeding probability increases with precipitation over the breeding season (Cayuela et al. 2014, 2017) and that adults focus their breeding effort on ponds with limited water level variation (Tournier et al. 2017). In our study system, individuals reproducing in variable sites had a higher propensity to skip breeding, likely because water level in ponds are often too low for successful tadpole development. In these sites, the proportion of individuals skipping a breeding opportunity was more variable over time than in stable ones, probably because individuals adjusted their breeding decisions according to pond water level and hydroperiod. This interpretation is in line with the observation that breeding population size of amphibians is often determined by rainfall, particularly in species breeding in temporary ponds (Pechmann et al. 1991). Our results show that these differences in breeding effort were associated with changes in the mean and the temporal variance of recruitment: both were higher in variable sites than in stable ones. A high synchronization of breeding probability with pond water levels likely increases the variance of offspring production over time. In variable sites, less regular reproduction probably results in higher variance of juvenile production, which may translate into more temporally variable adult recruitment into the adult population. Despite this broader variation, our findings show that mean recruitment was higher in variable sites than in stable ones. This could be explained by local hydrological conditions: adults may only reproduce during very wet years, which results in a higher breeding success when reproduction is possible. In addition, variation in environmental conditions prevailing during larval growth in the two types of sites could also explain this pattern. Both environmental conditions and biotic interactions are well known to differ between stable (closer to the permanent end of the hydroperiod gradient) and variable (more at the temporary end of the hydroperiod gradient) ponds (e.g., food levels, competition, predation; Wilbur 1980, 1987, Wellborn et al. 1996) and these conditions and interactions are known to affect age and size at metamorphosis (Smith 1983, Wilbur 1987, Newman 1992), which then strongly influence post-metamorphic development, survival, and age at maturity (Alvarez & Nicieza 2002, Van Allen et al. 2010, Schmidt et al. 2012).

### Stability and variability of breeding sites and the co-occurrence of contrasting life histories

Our results confirm theoretical predictions (Stearns 1976, Bulmer 1985, Tuljapurkar et al. 2009, Tuljapurkar 2013) about the relationships between life-history characteristics and environmental stochasticity. In populations experiencing highly stochastic breeding sites, lifespan was two times higher than in populations with less stochastic reproduction sites. This longer lifespan likely translates into a higher level of iteroparity, under the condition that sexual maturity is not delayed in these populations. Yet the slowing down of the life history was not associated with a lower temporal variance of adult survival, which suggests that a longer lifespan does not result from astrengthening of the canalization in adult survival in response to environmental variation. Rather, it more likely results from a trade-off between current reproduction and survival (Williams 1966, Stearns 1992, Ruf et al. 2006). By skipping breeding frequently in response to conditions unfavorable to reproduction, individuals may be able to save energy and/or to reduce mortality risks associated with reproductive activities, which extends toads’ lifespan.

Higher survival and recruitment rates should inevitably lead to a higher intrinsic growth rate in populations reproducing in variable sites. Because of density dependence, these populations should emit dispersers that would immigrate into neighboring populations, resulting in source-sink dynamics (Pulliam 1988). In our study system, dispersal occurs between populations but we detected too few dispersal events – as commonly assumed in the classical Levins’ metapopulation model (Harrison 1991) – to integrate this process in our capture-recapture models (Brandt 2015). After many generations, in a system at equilibrium, one should expect that the ‘variable’ strategy will dominate in the metapopulation if (1) it has a heritable genetic basis, if (2) individuals with the ‘variable’ phenotype do not experience a high fitness cost when they immigrate to stable sites (i.e. counter-selection through mortality or reproduction loss; Bonte et al. 2012), and if (3) individuals with the ‘variable’ phenotype do not immigrate preferentially into variable sites (i.e. ‘habitat matching’; Edelaar et al. 2008). By contrast, it is expected that the two strategies will coexist in the metapopulation if at least one of these assumptions is false. In particular, it is possible that adaptative phenotypic plasticity, rather than adaptation based on heritable genetic variation, allows toads to cope with stochastic variation of pond water levels (Newman 1992). These conjectures should be tested in future studies. It would also be interesting to test whether human habitat modification contributed to the divergence of life histories that we describe here. Variable ponds were ponds with natural variation in hydroperiod whereas ponds in stable sites were primarily man-made. It could therefore be that the stabilization of hydroperiods affects the life history of species. Since human activity affected life histories in other species (Alberto et al. 2017), it might happen in amphibians as well.

In conclusion, we show that the relationships between life-history variation and environmental stochasticity detected between species can also be found at the intraspecific level within a metapopulation. Thus, our results indicate similarities in the macroevolutionary and microevolutionary processes shaping life-history evolution.

## Acknowledgments

We thank the Bundesamt für Umwelt for funding the study (Forschungsvertrag 13.0007.PJ/M214-0617 to BRS). We also thank Martin Jordan and Daniel Hasen who collected the first two years of the mark-recapture data. In addition, we thank the Laboratorium der Urkantone for the permits to conduct the mark-recapture study (permit nos. 11-025 and 12-009). Finally, we warmly thank all the volunteers who contributed to the fieldwork, and Sarah Bänziger and Matteo Brezzi for assistance constructing capture histories. We thank Martin Wehrle (Tierpark Goldau) for providing accommodation.

## Supplementary material 1

**Table S1.**
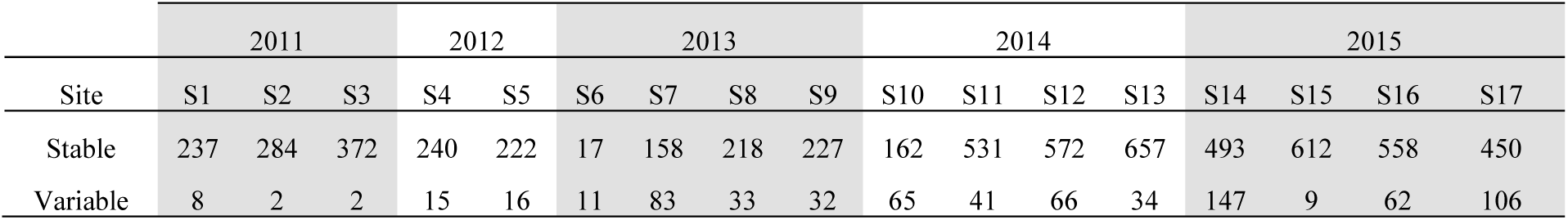
Number of individuals captured at each capture sessions over the 5-years period in the stable and variable sites.

**Table S1.** Survival analyses: model selection procedure. We show the 10 best-supported models based on their AICc rank (the complete selection procedure is provided in Supplementary material 2). The models include four parameters: φ **=** survival, γ_*intra*_ = intra-annual skipping breeding probability, γ_*inter*_ = inter-annual skipping breeding probability, *p* = recapture probability. k = number of model parameters, Deviance = residual deviance, ΔAICc = difference of AICc points between the model and the best-supported model.

| Model | k | Deviance | $\Delta AICc$ | w |
| --- | --- | --- | --- | --- |
| $\phi(\text{Site}), Y_{intra}(\text{Site}), Y_{inter}(\text{Site}), p(\text{Het+Year+Site})$ | 14 | 16772.94 | 0.00 | 0.57 |
| $\phi(\text{Site}), Y_{intra}(\text{Site}), Y_{inter}(\text{Site}), p(\text{Het+Year})$ | 13 | 16775.58 | 0.63 | 0.42 |
| $\phi(.), Y_{intra}(\text{Site}), Y_{inter}(\text{Site}), p(\text{Het+Year+Site})$ | 13 | 16783.35 | 8.40 | 0.01 |
| $\phi(\text{Site}), Y_{intra}(.), Y_{inter}(.), p(\text{Het+Year+Site})$ | 13 | 16786.71 | 11.76 | 0.00 |
| $\phi(.), Y_{intra}(\text{Site}), Y_{inter}(\text{Site}), p(\text{Het+Year})$ | 12 | 16790.66 | 13.70 | 0.00 |
| $\phi(\text{Site}), Y_{intra}(.), Y_{inter}(.), p(\text{Het+Year})$ | 12 | 16791.41 | 14.45 | 0.00 |
| $\phi(.), Y_{intra}(.), Y_{inter}(.), p(\text{Het+Year})$ | 11 | 16797.80 | 18.83 | 0.00 |
| $\phi(.), Y_{intra}(.), Y_{inter}(.), p(\text{Het+Year+Site})$ | 12 | 16797.70 | 20.74 | 0.00 |
| $\phi(\text{Site}), Y_{intra}(\text{Site}), Y_{inter}(\text{Site}), p(\text{Year+Site})$ | 12 | 17041.53 | 264.57 | 0.00 |
| $\phi(.), Y_{intra}(\text{Site}), Y_{inter}(\text{Site}), p(\text{Year+Site})$ | 11 | 17060.77 | 281.80 | 0.00 |
| $\phi(\text{Site}), Y_{intra}(\text{Site}), Y_{inter}(\text{Site}), p(\text{Year})$ | 11 | 17098.45 | 319.48 | 0.00 |
| $\phi(\text{Site}), Y_{intra}(.), Y_{inter}(.), p(\text{Year+Site})$ | 11 | 17100.00 | 321.04 | 0.00 |
| $\phi(.), Y_{intra}(\text{Site}), Y_{inter}(\text{Site}), p(\text{Year})$ | 10 | 17116.59 | 335.61 | 0.00 |
| $\phi(.), Y_{intra}(.), Y_{inter}(.), p(\text{Year+Site})$ | 10 | 17117.32 | 336.35 | 0.00 |
| $\phi(\text{Site}), Y_{\text{intra}}(\text{Site}), Y_{\text{inter}}(\text{Site}), p(\text{Het+Site})$ | 10 | 17118.98 | 338.01 | 0.00 |
| $\phi(\text{Site}), Y_{\text{intra}}(\cdot), Y_{\text{inter}}(\cdot), p(\text{Year})$ | 9 | 17121.57 | 338.59 | 0.00 |
| $\phi(\cdot), Y_{\text{intra}}(\cdot), Y_{\text{inter}}(\cdot), p(\text{Year})$ | 8 | 17127.19 | 342.21 | 0.00 |
| $\phi(\text{Site}), Y_{\text{intra}}(\cdot), Y_{\text{inter}}(\cdot), p(\text{Het+Site})$ | 9 | 17148.90 | 365.92 | 0.00 |
| $\phi(\cdot), Y_{\text{intra}}(\text{Site}), Y_{\text{inter}}(\text{Site}), p(\text{Het+Site})$ | 9 | 17150.49 | 367.51 | 0.00 |
| $\phi(\text{Site}), Y_{\text{intra}}(\text{Site}), Y_{\text{inter}}(\text{Site}), p(\text{Het})$ | 9 | 17153.43 | 370.45 | 0.00 |
| $\phi(\cdot), Y_{\text{intra}}(\text{Site}), Y_{\text{inter}}(\text{Site}), p(\text{Het})$ | 8 | 17158.06 | 373.08 | 0.00 |
| $\phi(\text{Site}), Y_{\text{intra}}(\cdot), Y_{\text{inter}}(\cdot), p(\text{Het})$ | 7 | 17171.05 | 384.06 | 0.00 |
| $\phi(\cdot), Y_{\text{intra}}(\cdot), Y_{\text{inter}}(\cdot), p(\text{Het+Site})$ | 8 | 17183.53 | 398.55 | 0.00 |
| $\phi(\cdot), Y_{\text{intra}}(\cdot), Y_{\text{inter}}(\cdot), p(\text{Het})$ | 6 | 17188.89 | 399.90 | 0.00 |
| $\phi(\text{Site}), Y_{\text{intra}}(\text{Site}), Y_{\text{inter}}(\text{Site}), p(\text{Site})$ | 7 | 17441.48 | 654.49 | 0.00 |
| $\phi(\cdot), Y_{\text{intra}}(\text{Site}), Y_{\text{inter}}(\text{Site}), p(\text{Site})$ | 6 | 17466.48 | 677.49 | 0.00 |
| $\phi(\text{Site}), Y_{\text{intra}}(\cdot), Y_{\text{inter}}(\cdot), p(\text{Site})$ | 6 | 17466.74 | 677.75 | 0.00 |
| $\phi(\text{Site}), Y_{\text{intra}}(\text{Site}), Y_{\text{inter}}(\text{Site}), p(\cdot)$ | 6 | 17467.99 | 679.00 | 0.00 |
| $\phi(\text{Site}), Y_{\text{intra}}(\cdot), Y_{\text{inter}}(\cdot), p(\cdot)$ | 5 | 17492.62 | 701.62 | 0.00 |
| $\phi(\cdot), Y_{\text{intra}}(\cdot), Y_{\text{inter}}(\cdot), p(\cdot)$ | 5 | 17494.33 | 703.34 | 0.00 |
| $\phi(\cdot), Y_{\text{intra}}(\cdot), Y_{\text{inter}}(\cdot), p(\text{Site})$ | 6 | 17493.62 | 704.62 | 0.00 |
| $\phi(\cdot), Y_{\text{intra}}(\text{Site}), Y_{\text{inter}}(\text{Site}), p(\cdot)$ | 6 | 17495.92 | 706.93 | 0.00 |

**Table S2.** Recruitment analyses: model selection procedure. We show the 10 best-supported models based on their AICc rank (the complete selection procedure is provided in Supplementary material 2). The models include four parameters: Ψ **=** recruitment, γ_*intra*_ = intra-annual skipping breeding probability, γ_*inter*_ = inter-annual skipping breeding probability, *p* = recapture probability. k = number of model parameters, Deviance = residual deviance, ΔAICc = difference of AICc points between the model and the best-supported model.

| Model | k | Deviance | $\Delta\text{AICc}$ | w |
| --- | --- | --- | --- | --- |
| $\psi(\text{Site}), Y_{\text{intra}}(\text{Site}), Y_{\text{inter}}(\text{Site}), p(\text{Het+Year+Site})$ | 14 | 18098.58 | 0.00 | 0.89 |
| $\psi(\cdot), Y_{\text{intra}}(\text{Site}), Y_{\text{inter}}(\text{Site}), p(\text{Het+Year+Site})$ | 13 | 18104.85 | 4.25 | 0.11 |
| $\psi(\cdot), Y_{\text{intra}}(\text{Site}), Y_{\text{inter}}(\text{Site}), p(\text{Het+Year})$ | 12 | 18113.02 | 10.42 | 0.00 |
| $\psi(\text{Site}), Y_{\text{intra}}(\text{Site}), Y_{\text{inter}}(\text{Site}), p(\text{Het+Year})$ | 13 | 18112.79 | 12.20 | 0.00 |
| $\psi(\cdot), Y_{\text{intra}}(\cdot), Y_{\text{inter}}(\cdot), p(\text{Het+Year+Site})$ | 12 | 18139.94 | 37.34 | 0.00 |
| $\psi(\text{Site}), Y_{\text{intra}}(\cdot), Y_{\text{inter}}(\cdot), p(\text{Het+Year+Site})$ | 13 | 18138.16 | 37.57 | 0.00 |
| $\psi(\text{Site}), Y_{\text{intra}}(\cdot), Y_{\text{inter}}(\cdot), p(\text{Het+Year})$ | 12 | 18157.17 | 54.57 | 0.00 |
| $\psi(\cdot), Y_{intra}(\cdot), Y_{inter}(\cdot), p(\text{Het}+\text{Year})$ | 11 | 18161.29 | 56.68 | 0.00 |
| $\psi(\text{Site}), Y_{intra}(\text{Site}), Y_{inter}(\text{Site}), p(\text{Year}+\text{Site})$ | 12 | 18258.68 | 156.08 | 0.00 |
| $\psi(\cdot), Y_{intra}(\text{Site}), Y_{inter}(\text{Site}), p(\text{Year}+\text{Site})$ | 11 | 18265.83 | 161.23 | 0.00 |
| $\psi(\cdot), Y_{intra}(\text{Site}), Y_{inter}(\text{Site}), p(\text{Year})$ | 10 | 18298.20 | 191.58 | 0.00 |
| $\psi(\text{Site}), Y_{intra}(\text{Site}), Y_{inter}(\text{Site}), p(\text{Year})$ | 11 | 18298.09 | 193.48 | 0.00 |
| $\psi(\cdot), Y_{intra}(\cdot), Y_{inter}(\cdot), p(\text{Year}+\text{Site})$ | 10 | 18325.61 | 218.99 | 0.00 |
| $\psi(\text{Site}), Y_{intra}(\cdot), Y_{inter}(\cdot), p(\text{Year}+\text{Site})$ | 11 | 18325.38 | 220.77 | 0.00 |
| $\psi(\text{Site}), Y_{intra}(\cdot), Y_{inter}(\cdot), p(\text{Year})$ | 9 | 18340.46 | 231.84 | 0.00 |
| $\psi(\cdot), Y_{intra}(\cdot), Y_{inter}(\cdot), p(\text{Year})$ | 8 | 18346.66 | 236.04 | 0.00 |
| $\psi(\cdot), Y_{intra}(\text{Site}), Y_{inter}(\text{Site}), p(\text{Het})$ | 8 | 18983.98 | 873.35 | 0.00 |
| $\psi(\text{Site}), Y_{intra}(\text{Site}), Y_{inter}(\text{Site}), p(\text{Het})$ | 9 | 18983.66 | 875.04 | 0.00 |
| $\psi(\cdot), Y_{intra}(\text{Site}), Y_{inter}(\text{Site}), p(\text{Het}+\text{Site})$ | 9 | 18983.71 | 875.09 | 0.00 |
| $\psi(\cdot), Y_{intra}(\cdot), Y_{inter}(\cdot), p(\text{Het}+\text{Site})$ | 8 | 18986.33 | 875.71 | 0.00 |
| $\psi(\cdot), Y_{intra}(\cdot), Y_{inter}(\cdot), p(\text{Het})$ | 8 | 18986.33 | 875.71 | 0.00 |
| $\psi(\text{Site}), Y_{intra}(\text{Site}), Y_{inter}(\text{Site}), p(\text{Het}+\text{Site})$ | 10 | 18983.54 | 876.93 | 0.00 |
| $\psi(\text{Site}), Y_{intra}(\cdot), Y_{inter}(\cdot), p(\text{Het}+\text{Site})$ | 9 | 18986.32 | 877.70 | 0.00 |
| $\psi(\text{Site}), Y_{intra}(\cdot), Y_{inter}(\cdot), p(\text{Het})$ | 9 | 18986.32 | 877.70 | 0.00 |
| $\psi(\cdot), Y_{intra}(\text{Site}), Y_{inter}(\text{Site}), p(\cdot)$ | 6 | 19128.23 | 1013.60 | 0.00 |
| $\psi(\cdot), Y_{intra}(\text{Site}), Y_{inter}(\text{Site}), p(\text{Site})$ | 6 | 19128.23 | 1013.60 | 0.00 |
| $\psi(\cdot), Y_{intra}(\cdot), Y_{inter}(\cdot), p(\text{Site})$ | 5 | 19130.59 | 1013.96 | 0.00 |
| $\psi(\text{Site}), Y_{intra}(\text{Site}), Y_{inter}(\text{Site}), p(\text{Site})$ | 7 | 19127.87 | 1015.25 | 0.00 |
| $\psi(\text{Site}), Y_{intra}(\text{Site}), Y_{inter}(\text{Site}), p(\cdot)$ | 7 | 19127.87 | 1015.25 | 0.00 |
| $\psi(\text{Site}), Y_{intra}(\cdot), Y_{inter}(\cdot), p(\text{Site})$ | 6 | 19130.43 | 1015.80 | 0.00 |
| $\psi(\text{Site}), Y_{intra}(\cdot), Y_{inter}(\cdot), p(\cdot)$ | 5 | 19144.18 | 1027.54 | 0.00 |
| $\psi(\cdot), Y_{intra}(\cdot), Y_{inter}(\cdot), p(\cdot)$ | 4 | 19151.04 | 1032.40 | 0.00 |

**Fig. S1.**
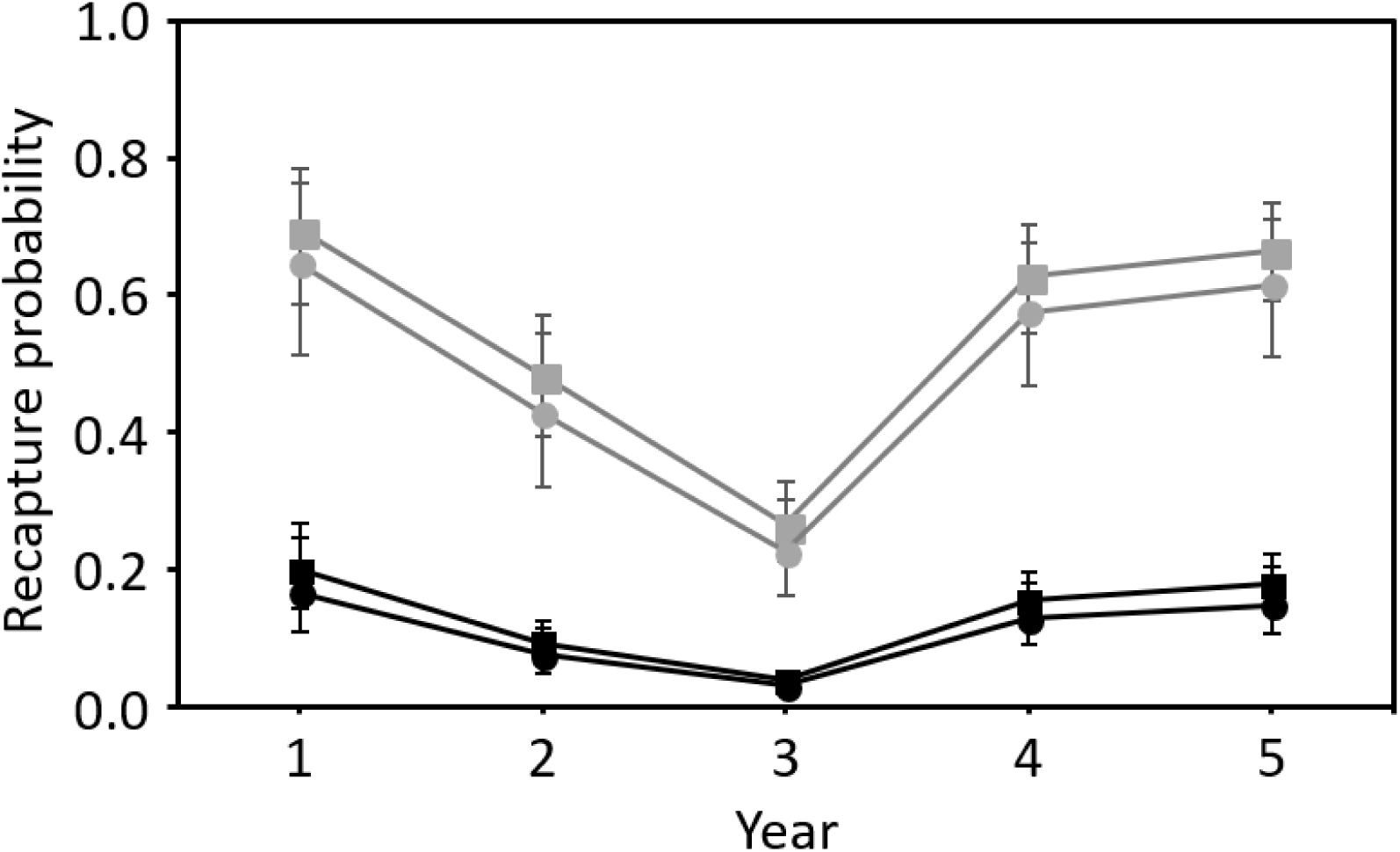
Model-averaged recapture probabilities extracted from survival models. The recapture probabilities estimated for the heterogeneity mixture 1 are shown in black, for heterogeneity mixture 2 in grey. The recapture probabilities in predictable sites are shown as squares, in predictable sites as circles.

